# Characterizing Primary transcriptional responses to short term heat shock in paired fraternal lymphoblastoid lines with and without Down syndrome

**DOI:** 10.1101/2023.01.17.524431

**Authors:** Joseph F. Cardiello, Jessica Westfall, Robin Dowell, Mary Ann Allen

**Affiliations:** BioFrontiers Institute,University of Colorado, Boulder, United States; Molecular, Cellular and Developmental Biology, University of Colorado, Boulder, United States; Linda Crnic Institute, University of Colorado, Denver, United States; BioFrontiers-Crnic Boulder Branch,University of Colorado, Boulder, United States; Molecular Medicine and Gene Therapy, Wallenberg Centre for Molecular Medicine, Lund Stem Cell Center, Lund University, Lund, Sweden

## Abstract

Heat shock stress induces genome wide changes in transcription regulation, activating a coordinated cellular response to enable survival. Using publicly available transcriptomic and proteomic data sets comparing individuals with and without trisomy 21, we noticed many heat shock genes are up-regulated in blood samples from individuals with trisomy 21. Yet no major heat shock response regulating transcription factor is encoded on chromosome 21, leaving it unclear why trisomy 21 itself would cause a heat shock response, or how it would impact the ability of blood cells to subsequently respond when faced with heat shock stress. To explore these issues in a context independent of any trisomy 21 associated co-morbidities or developmental differences, we characterized the response to heat shock of two lymphoblastoid cell lines derived from brothers with and without trisomy 21. To carefully compare the chromatin state and the transcription status of these cell lines, we measured nascent transcription, chromatin accessibility, and single cell transcript levels in the lymphoblastoid cell lines before and after acute heat shock treatment. The trisomy 21 cells displayed a more robust heat shock response after just one hour at 42°C than the matched disomic cells. We suggest multiple potential mechanisms for this increased heat shock response in lymphoblastoid cells with trisomy 21 including the possibility that cells with trisomy 21 may exist in a hyper-reactive state due to chronic stresses. Whatever the mechanism, abnormal heat shock response in individuals with Down syndrome may hobble immune responses during fever and contribute to health problems in these individuals.

## Introduction

Trisomy 21 is caused by an extra copy of chromosome 21. For the most part, it is unclear how this extra copy of 1% of the genome leads to phenotypes associated with trisomy 21. Trisomy 21 cells have been demonstrated to show an increased reaction to key cellular perturbations. For instance, trisomy 21 cells show an increased interferon response relative to typical cells which is likely driven by overexpression of four interferon receptor genes encoded on chromosome 21(***Sullivan et al. (2016***); ***Araya et al. (2019***); ***Powers et al. (2019***); ***Waugh et al. (2019***)). Similarly, trisomic cells show signs of an elevated oxidative stress response, which may tie into the chromosome 21 encoded, oxidative stress responsive NRF2 or SOD1 genes (***Lanzillotta and Di Domenico (2021***); ***Zhu et al. (2019***); ***Pagano and Castello (2012***)). Despite the clear ties between these chromosome 21 located genes and the unusual cellular responses, it is unclear if the increased response to these two perturbations is driven solely by regulatory genes encoded on chromosome 21 or if other factors contribute such as a general trisomic stress response, or a trisomy 21 derived chronic stress. Moreover, it is unclear how trisomy 21 cells respond to perturbations that do not have primary regulators on chromosome 21.

Heat shock is a potentially lethal stress and therefore it activates a variety of cellular response processes including the unfolded protein response. While three genes encoded on chromosome 21 are heat shock activated (HSF2BP, DNAJC28, HSPA13), none of these heat shock genes are known to be upstream regulators of the heat shock response. The major regulator of heat shock, HSF1, is a transcription factor located on chromosome 8. HSF1 is ubiquitously expressed, but its transcription factor activity is highly regulated through post translational modifications, nuclear import, and protein interactions (***Anckar and Sistonen (2011)***; ***Björk and Sistonen (2010)***; ***Voellmy (2004)***; ***Vihervaara and Sistonen (2014)***). Following heat shock there is an increase in HSF1 DNA binding and HSF1 activity (reviewed in ***Anckar and Sistonen (2011)***; ***Huang et al. (2018***)). Activation of HSF1 results in genome wide transcription changes including activation of the production of heat shock proteins, and repression of thousands of genes (***Ray et al. (2019***); ***Mahat et al. (2016***)).

We used heat shock to investigate whether trisomy 21 impacts the ability of cells to mount a robust stress response in which the master regulators do not reside on chromosome 21. Previous studies provide mixed messages about how aneuploid cells respond to heat shock. A study by Beach et al found that aneuploidy in yeast leads to increased cell to cell variation in response to heat shock (***Beach et al. (2017***)). Another study found that human fibroblasts with trisomy 21 failed to properly activate the expression of a couple of key heat shock proteins after heat shock (***Aivazidis et al. (2017***)). In published untreated clinical blood samples, we found an irregular elevation of heat shock target genes in individuals with trisomy 21 ***Figure 2***. Therefore, to more directly assess whether trisomy 21 blood cells properly respond to heat shock, we examined the primary effects of heat shock on lymphoblastoid cell lines from two brothers. We found by multiple omics assays that after a short, mild heat shock stress, the trisomy 21 lymphoblastoid cell line activates primary HSF1 regulated transcriptional responses more robustly than the diploid cells.

**Figure 1.**
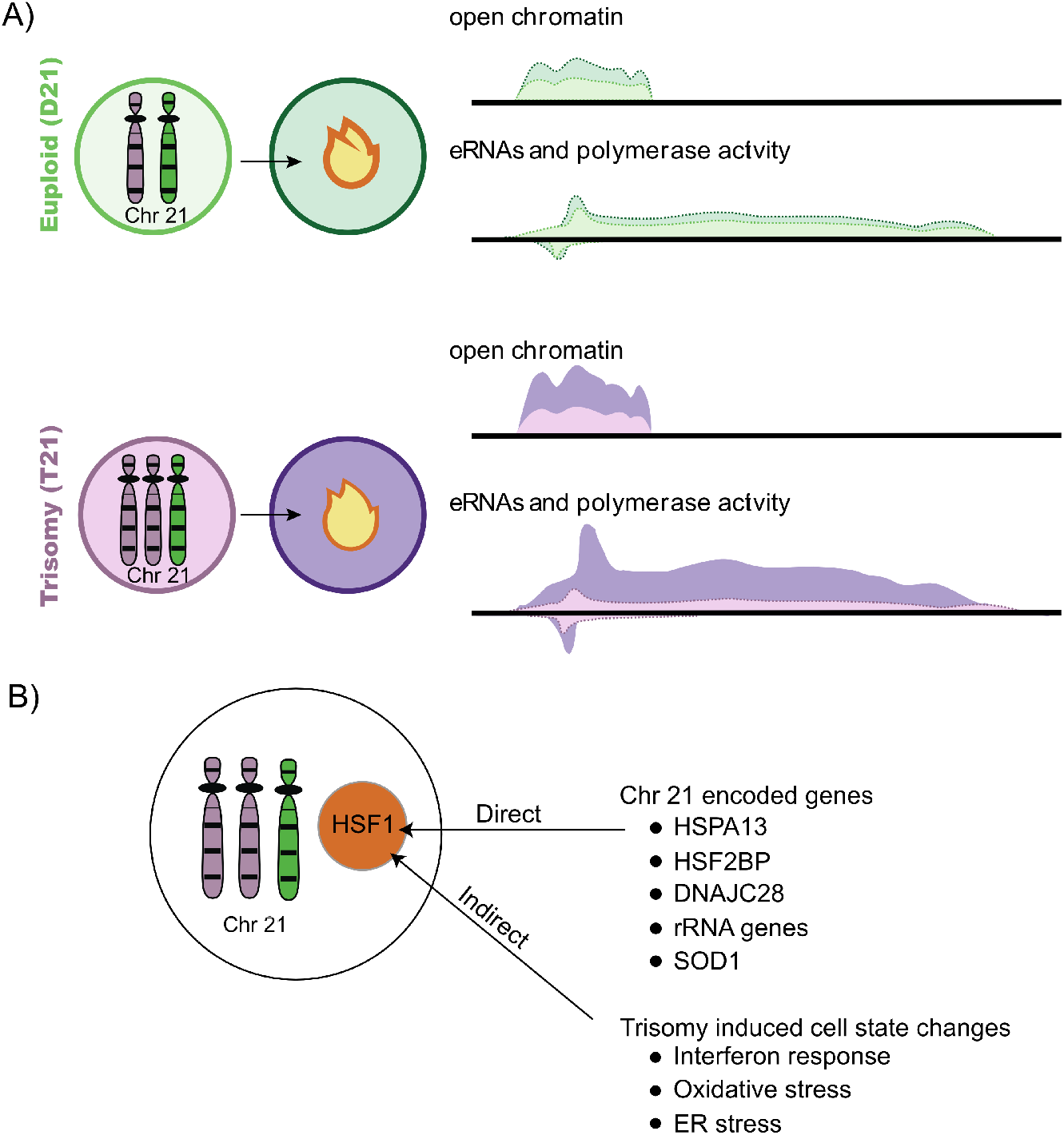
Graphical abstract. A: Cells with trisomy 21 have increased acute heat shock activated HSF1 transcription factor function as determined by both PRO-seq and ATAC-seq analysis. B: Several trisomy 21 cellular changes may contribute to an increased response to heat shock.

**Figure 2.**
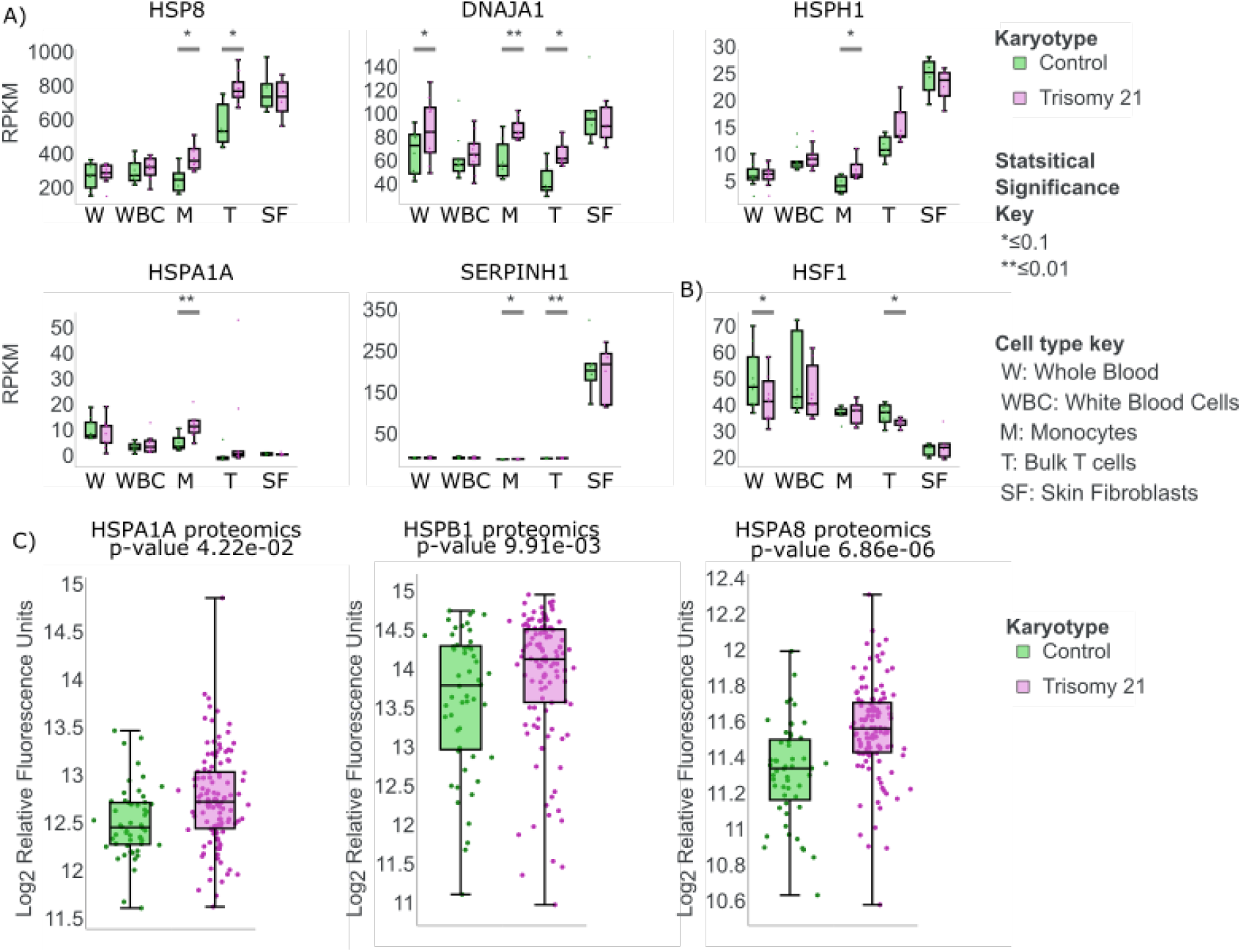
Individuals with trisomy 21 have elevated levels of some heat shock regulated genes under normal conditions. All Data from the Human Trisome project ***Linda Crnic Institute (2021)***. A: Several heat shock genes (HSPA8, DNAJA1, HSPH1, HSPA1A, SERPINH1) are differentially expressed (RNA-seq) in multiple blood cell lineages in individuals with trisomy 21 (purple) compared to disomic controls (green). Multiple clinical samples shown: whole blood (W), white blood cells (WBC), monocytes (M), bulk T cells (T), and skin fibroblasts (SF). Significance key: * ≤ 0.1, ** ≤ 0.01. B: HSF1 transcript levels in the same blood cell lineages. C: Clinical blood sample proteomics data (***Sullivan et al. (2017***)) shows elevated levels for some heat shock induced genes in plasma from people with trisomy 21. **Figure 2–Figure supplement 1**. Heat shock genes altered in transcript or protein levels.

## Results

### Individuals with trisomy 21 have elevated levels of genes related to heat shock in some blood cell lineages

To determine whether heat shock response is influenced by trisomy 21, we first examined publicly available clinical data. Several projects have collected unperturbed RNA-seq data from clinical samples of multiple blood and skin cell types in individuals with and without trisomy 21, including the Human Trisome Project***Linda Crnic Institute (2021)***. In the Human Trisome Project samples (***Linda Crnic Institute (2021)***; ***Sullivan et al. (2016***); ***Araya et al. (2019***); ***Powers et al. (2019***); ***Waugh et al. (2019***), data accessed May 23rd, 2022), we noted that the RNA levels for several heat shock regulated genes were higher in individuals with trisomy 21, particularly in T cells and monocytes (see ***Figure 2***A). Specifically, transcript levels for the heat shock regulated genes HSPA8, DNAJA1, HSPH1, HSPA1A, and SERPINH1 are elevated in trisomy 21 groups compared to clinical controls. Furthermore, the transcript levels for HSF1, the master heat shock regulating transcription factor, appears changed in some trisomy 21 cell types (see ***Figure 2***B). Published proteomic data further confirmed that HSPA1A, HSPA8, and DNAJB1 protein levels are elevated in blood samples from individuals with trisomy 21 (***Sullivan et al. (2017***)). Importantly, transcript and proteomic levels were not increased consistently for all HSF1 regulated genes in all blood cell types (***Figure 2–Figure Supplement 1***). Thus the clinical data present a perplexing picture of how trisomy 21 influences the expression of heat shock related genes.

### Greater heat shock induced increase in chromatin accessibility at HSF1 sites in trisomic cells

To investigate the trisomic heat shock response outside of clinical complexities, we set out to characterize the acute heat shock response in paired cell lines with and without trisomy 21 (***Figure 3***A). To determine how chromatin accessibility changes as blood cells respond to heat shock, age and gender matched lymphoblastoid cells derived from two brothers were assayed for transposase-accessible chromatin (ATAC-seq) under control conditions 37 °C, and mild heat shock treatment, e.g. 1 hour at 42 °C (See ***Figure 3***A for design). We used lymphoblastoid cells because they are a readily available blood like cell type. We used lymphoblastoid cells because they are a readily available blood like cell type. A short time point (1 hour) was chosen to focus on the primary response to heat shock, as this avoids most secondary or downstream effects that arise from cellular feedback mechanisms.

**Figure 3.**
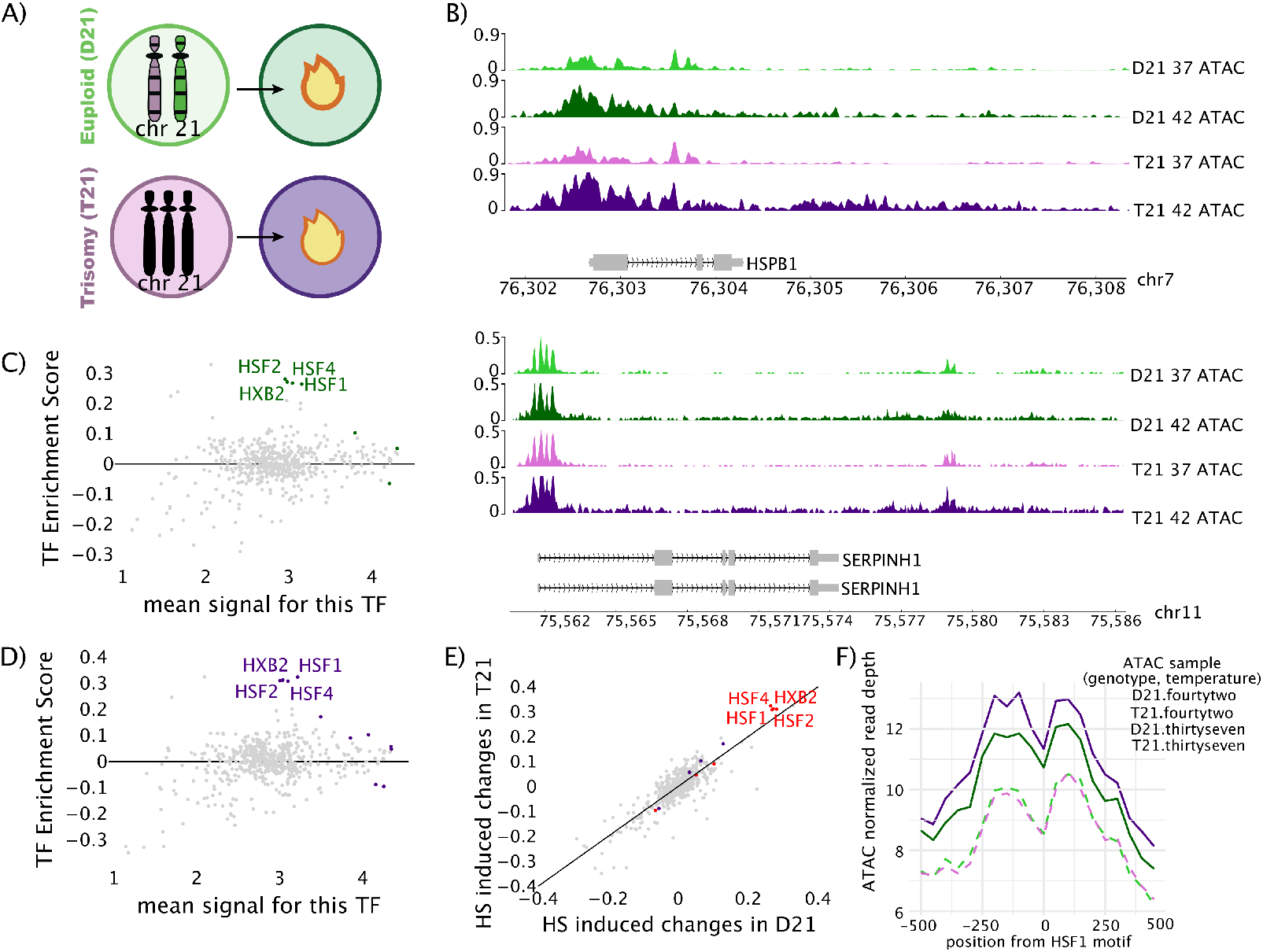
After acute heat shock, cells with trisomy 21 have increased chromatin accessibility near heat shock response elements compared to disomic controls. A: Conceptual diagram of the conditions analyzed. Disomic cells (green) and trisomic cells (purple) at control (37°, light color) or mild heat shock (42°, dark color). B: ATAC-seq data traces of two heat shock regulated genes (HSPB1, SERPHIN1). All traces for an individual gene have the same y-axis scale. C: MA plot of heat shock induced changes in transcription factor activity in the disomic cell line based on TFEA (Transcription Factor Enrichment Analysis) analysis ***Rubin et al. (2021***) of ATAC-seq data. Grey: non-siginficant TFs, colored: GC-corrected P-adjusted value of p<1×10^−10^. D: MA plot of heat shock induced changes in transcription factor activity in the trisomic cell line based on TFEA analysis ***Rubin et al. (2021***) of ATAC-seq data. Grey: non-siginficant TFs, colored: GC-corrected P-adjusted value of p<1×10^−10^. E: Scatter plot comparing heat shock induced changes in transcription factor activity (E-values from TFEA) between disomic cells (X-axis) and trisomic cells (y-axis). Red: significant in both comparisons, Purple: Significant in trisomy only, Green: Significant in disomy only. F: Averaged signal of ATAC-seq data for 1 kilobase region around active HSF1 motifs. **Figure 3–Figure supplement 1**. Extended heatmaps of ATAC-seq and PRO-seq signal over HSF1 sites

We first confirmed the heat shock response was observable under these conditions in both cell lines by manual inspection of several well-known heat shock genes including HSPB1 and SERPINH1 (***Figure 3***B). These genes showed the expected response, i.e. opening of the chromatin at the promoter region in heat shock compared to control. We called peaks in this data using HMMRATAC and determined a list of peaks that were differentially expressed after heat shock in the trisomic cells and the disomic cells. All genes within 25 kilobases of a differential ATAC-seq peak were used for GO analysis. GO analysis showed the heatshock pathway was strongly activated in both the trisomy 21 and disomic samples. Interestingly, more peaks were called as differently expressed between the control and heat shocked conditions in the trisomic samples than the disomic samples.

We next sought to infer changes in transcription factor activity in response to heat shock in an unbiased fashion for both cell lines. To this end, we employed transcription factor enrichment analysis (TFEA)***Rubin et al. (2021***) on HMMRATAC***Tarbell and Liu (2019)*** called peaks to determine which transcription factor motifs co-associate with observed changes in chromatin accessibility genome wide. TFEA calculates a Escore, or enrichment score, which measures the motif occurrence in regions of interest that have altered accessibility/transcription. For example, if heat shock causes a transcription factor to bind its motif, open chromatin and activates transcription nearby (like HSF1), the E-score for that TF will be positive and high. On the other hand, if a transcription factor is binding to its motif, opening chromatin and activating transcription at 37 °C and after heat shock that TF leaves DNA then the E-score for that will be negative. Note, transcription factors that repress transcription will show the opposite Escore pattern in PRO-seq. In our data, in both the disomic and trisomic cell lines, TFEA infers that the transcription factors HSF1, HSF2, HSF4, and HXB2 were robustly induced by heat shock (***Figure 3***C, ***Figure 3***D, ***Figure 3–Figure Supplement 1B***). In addition, we directly compared the ATAC-seq TFEA inferred heat shock induced TF changes between the two cell lines and found that the same TFs were significantly upregulated in activity, though the activation was slightly more robust in the trisomic cells after this short, mild heat shock treatment (red TFs ***Figure 3***E).

Therefore, we next sought confirm the above result by characterizing changes in accessibility at known bound HSF1 motifs. To this end, we downloaded HSF1 ChIP-seq data from lymphoblastoid lines and filtered the data for binding sites with the HSF1 motifs (***Zheng et al. (2018***), ***Mei et al. (2016***), ***Consortium et al. (2012)***). We then graphed the ATAC-seq signal over the known HSF1 bound sites genome wide ***Figure 3–Figure Supplement 1A***). Upon heat shock, both cell lines show increases in accessibility at HSF1 sites, but the trisomic cell line has a more open ATAC-seq signal post heat shock (***Figure 3***F, ***Figure 3–Figure Supplement 1A***). Overall, our ATAC-seq data suggests that trisomic cells display a slightly elevated chromatin accessibility at HSF1 bound sites after heat shock, compared to disomic cells. This lead us to question whether the difference in chromatin was leading to concomitant changes in gene transcription in the trisomic cells upon heat shock.

### Trisomic cells with have increased transcription at HSF1 motifs after heat shock

To compare observed changes in chromatin accessibility to changes in transcription, we preformed precision run-on sequencing (PRO-seq) in the trisomic and disomic cells at the same time points (before and 1 hr HS) used for the ATAC-seq (Figure ***Figure 3***A for design). In both cell lines, heat shock genes such as HSPB1 and SERPINH1 were transcribed at higher levels after heat shock (***Figure 4***A, ***Figure 4–Figure Supplement 1A***). We used DESeq2 to assess differential gene transcription after heat shock in both samples (Fig ***Figure 4***B, ***Figure 4***C). Many genes showed a reduction in transcription in response to heat shock in both cell lines (Fig ***Figure 4***C, Fig ***Figure 4–Figure Supplement 1B***). The trisomic sample revealed more genes with significant changes in transcription in response to heat shock than the disomic sample (***Figure 4***B, ***Figure 4***C). Moreover, genes that were differentially transcribed in both samples showed a general trend of being induced to a greater extent in the trisomic cell line (Fig ***Figure 4–Figure Supplement 1B***, ***Figure 4–Figure Supplement 1C***).

**Figure 4.**
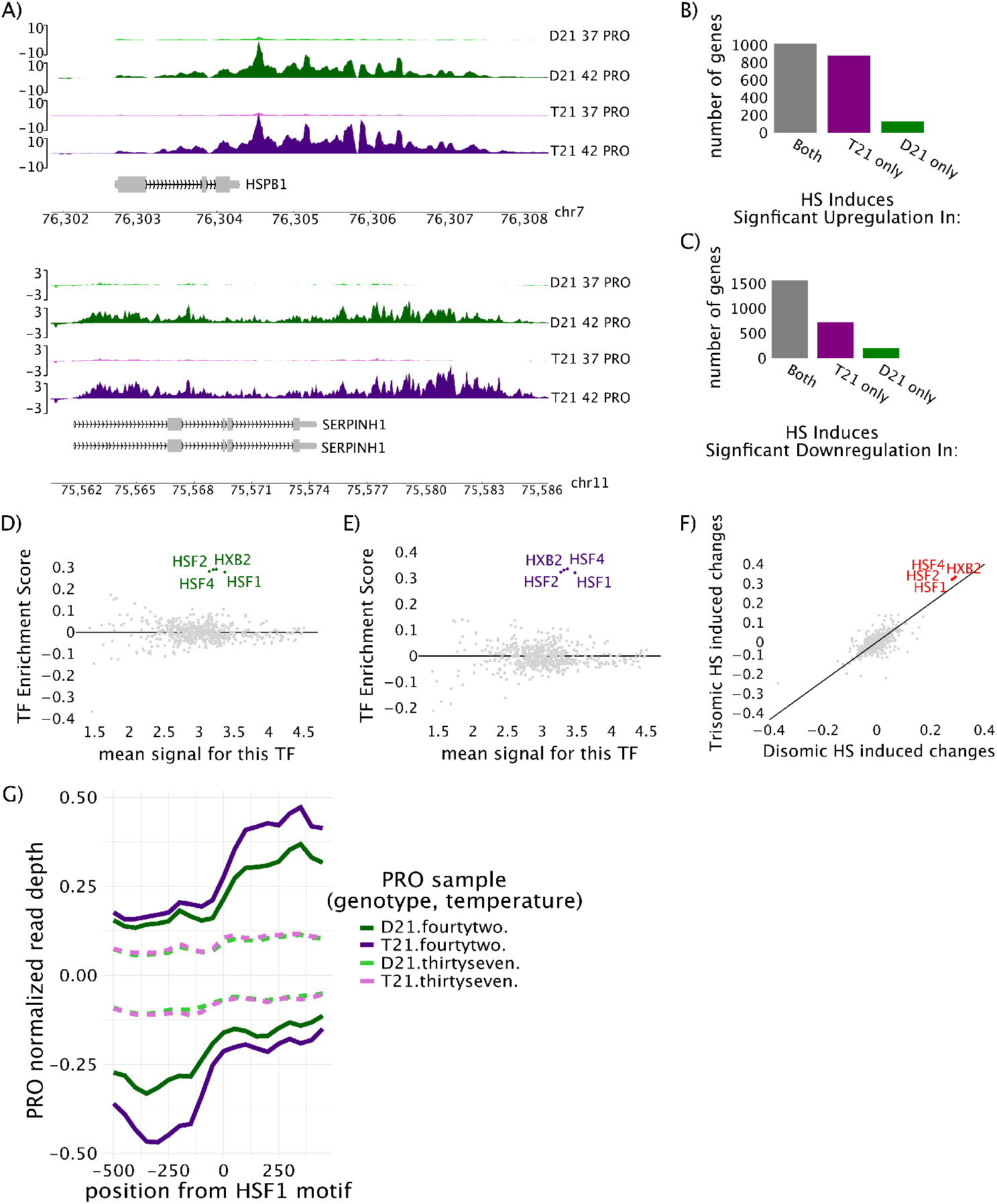
A mild heat shock treatment induces more robust transcriptional changes in the trisomic cell line compared to disomic control. A: Total read count corrected PRO-seq gene traces at two heat shock regulated genes (Hspb1, Serpinh1) in the four cell types/conditions. B: Number of genes with heat shock induced increases (by DESeq2) in gene transcription (PRO-seq) in one or both cell lines. C: Number of genes with heat shock induced repression (by DESeq2) of gene transcription (PRO-seq) identified in one or both cell lines. D: MA plot of heat shock induced changes in transcription factor activity in the disomic cell line, via TFEA analysis ***Rubin et al. (2021***). Grey: non-significant TFs, colored: significant at GC corrected p-adjusted value of p<1×10^−10^. E: MA plot of heat shock induced changes in transcription factor activity in the trisomic cell line, via TFEA analysis ***Rubin et al. (2021***). F: Scatter plot comparing TFEA derived GC-corrected E-score values for PRO-seq differences between disomic heat shock (X-axis) and trisomic heat shock (y-axis). Red: significant (p<1×10^−10^) in both comparisons G: Average metaplot of PRO-seq data surrounding (± 500 bp) lymphoblastoid-active HSF1 motifs (at zero). **Figure 4–Figure supplement 1**. Extended plots of PRO-seq gene transcription

To determine if HSF1 was the only TF with increased transcription associated with its motifs, we used TFEA to infer transcription factor activity changes based the PRO-seq data (independent of the changes in ATAC-seq). To this end, we used Tfit (Transcription fit) to identify all sites of bidirectional transcription within each PRO-seq data set (***Azofeifa and Dowell (2016)***). Regions of transcription were combined across conditions and replicates using muMerge. Consistent with the ATAC results, TFEA results on the PRO-seq data show a robust activation of HSF1, 2, and 4 in response to heat shock in both the disomic and trisomic cell line (Fig ***Figure 4***D, Fig ***Figure 4***E). Additionally, many TFs showed a subtle but non-significant reduction in activity in response to heat shock in both cell lines, consistent with widespread transcription repression in response to heat shock treatment (Fig ***Figure 4***D, Fig ***Figure 4***E). A direct comparison of the heat shock induced changes to TF activity inferred by PRO-seq signal revealed a higher relative activation of HSF TFs in the trisomic cell line compared to the disomic cell line (***Figure 4***F).

Since transcription factor binding sites co-occur with enhancer RNAs which are readily detected by the PRO-seq assay, we next examined nascent transcription at HSF1 binding sites in response to heat shock. We hypothesized that a more sensitive or robust HSF1 activation in the trisomic cells might explain the increased genome wide changes in chromatin accessibility and transcription in the trisomic cell line compared to the disomic line. Genome wide, heat shock led to an increase in PRO-seq signal at bound HSF1 motifs in both trisomic and disomic cell lines, confirming the activation of HSF1 motif adjacent eRNAs in both cell lines (Fig ***Figure 3–Figure Supplement 1C***, Fig ***Figure 4***G). Though PRO-seq levels began at similar levels in the two cell lines under control conditions, after just the one hour of mild heat shock treatment we noted a more robust transcriptional response in the trisomic cell line compared to the disomic cell line (Fig ***Figure 4***G). Collectively, both the PRO-seq and the ATAC-seq suggest that though the transcriptional response to heat shock is similar between the two cell lines, it is more robust in the trisomic cells after just one hour of mild heat shock.

### Single cell RNA sequencing confirms the increased heat shock response in trisomic cells is population wide rather than the result of outlier hyper-stressed or dying cells

The increase in heat shock response observed in trisomic cells could arise from either a small number of hyper responsive cells or a more consistent population wide effect. To address this question, we applied single cell RNA sequencing (scRNA-seq) to the same two cell lines at the same time points, namely before and 1 hr after heat shock. As a control, we first confirmed that the scRNA-seq protocol detected the expected higher quantity of chromosome 21 transcripts in the trisomy 21 sample. To this end, we plotted the depth normalized counts per cell for chromosome 21 encoded genes and found that these transcripts are present in higher quantities in the trisomic cell line than the disomic cell line (Fig ***Figure 5***A, Fig ***Figure 5***B). Additionally, we examined the transcript levels for known heat shock responsive genes and confirmed heat shock induced increases/decreases in the transcript level for these genes in both cell lines (Fig ***Figure 5***C, Fig ***Figure 5***D).

**Figure 5.**
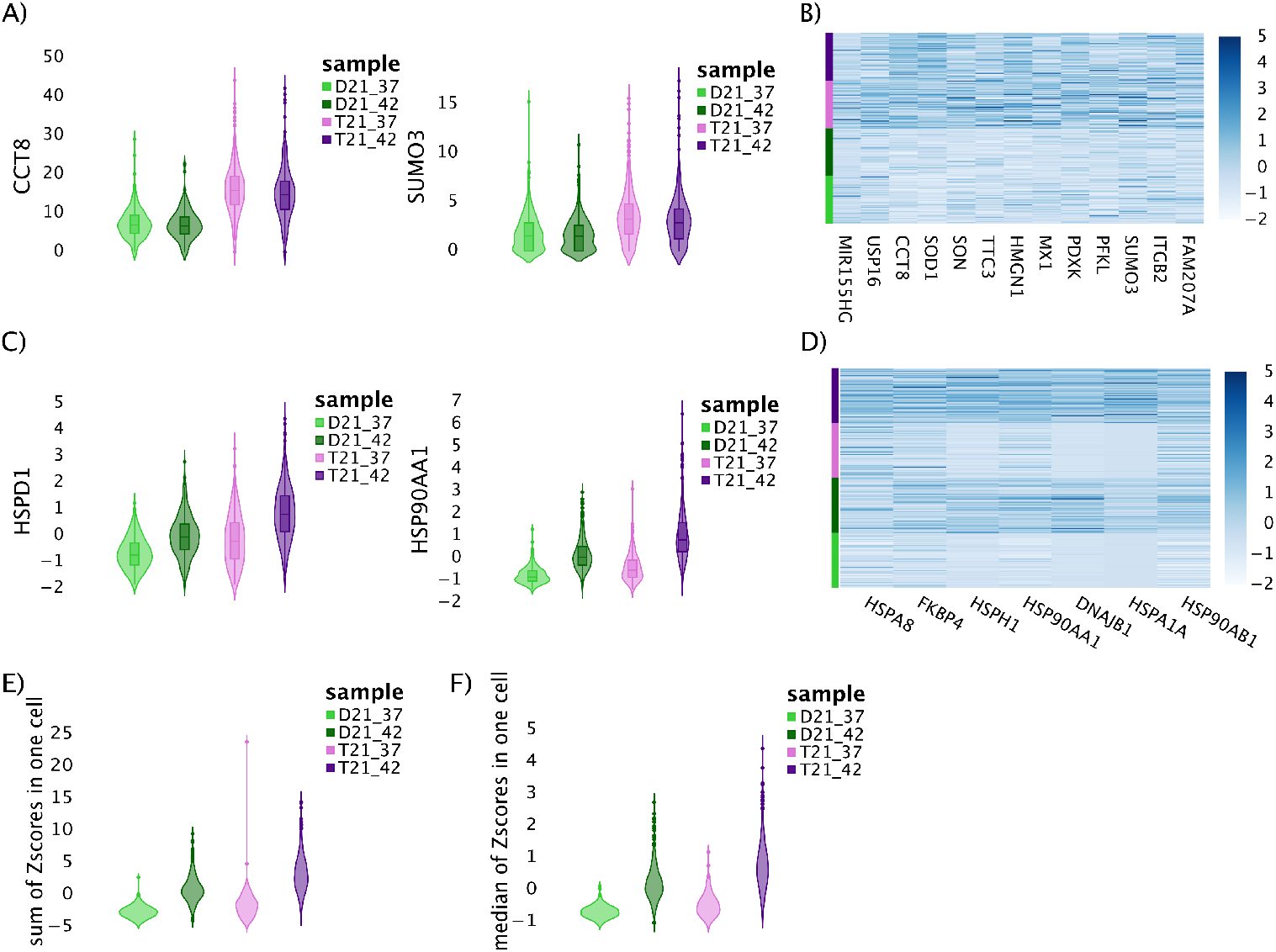
Single Cell RNA-seq indicates that the change in heat shock induced gene expression in trisomy 21 cells is population wide. A: Violin plots of percent of total normalized scRNA-seq gene counts per cell of two genes (CCT8, SUMO3) present on chromosome 21 in the control (light colors) and HS conditions (dark colors) of the disomic (green) and trisomic (purple) lymphoblastoid cell lines. B: Heatmap of the Z-scores of chromosome 21 genes showing a general up-regulation in expression of genes on chromosome 21. C: Violin plots of two heat shock responsive genes (HSPD1, HSP90AA1) not encoded on chromosome 21, the y-axis is the levels across more than 500 cells in each sample via single cell RNA-seq. D: Heatmap of the Z-scores of heat shock genes show a general up-regulation in expression of heat shock genes, rather than a few cells with extreme heat shock phenotypes. E: Violin plots showing of the sum of Z-scores of all heat shock genes for more than 500 cells in each sample. The genes showed must be present in at least 75% of cells. F: Violin plots showing the median Z-score of all heat shock genes for more than 500 cells in each sample. **Figure 5–Figure supplement 1**. Extended plots of scRNAseq

To address whether the observed increase in trisomy heat shock response was driven by a small number of outlier cells, we summed all Z-scores for all heat shock genes across individual cells. If a small population of cells had an usually strong heat shock response, those cells should be expected to have a sum that was an outlier, exceedingly higher than the other cells. Furthermore, the non-outlier cells would be expected to have sums roughly equal to that of the disomic cells. On the other hand, if the whole population of trisomic cells was showing a slightly more robust heat shock then the entire population of disomic cells, then the distribution of cell sums would be shifted. The results show that there is not a small population of super responsive heat shock cells in the trisomy sample. Instead the heat shock difference appears to be population wide: essentially all trisomic cells appear to have an increase in heat shock transcripts relative to the disomic cells (Fig ***Figure 5***E). The same result is obtained when using the median of Z-scores rather than the mean (Fig ***Figure 5***F). Although some cells respond more strongly than others to heat shock, the T21 cells do not show a different overall pattern than the D21 cells. This suggests that the increased heat shock response in trisomic cell lines is a population wide phenomenon, not the result of a small number of hyper responding cells.

## Discussion

In this study we found that the presence of a third copy of chromosome 21 did not disrupt the cellular ability to mount a heat shock response. Rather, we observed that the trisomic cells were surprisingly agile at changing gene expression in response to this mild perturbation and appeared to increase chromatin accessibility and transcription at HSF1 motifs more readily at this early time point, than the disomic control. Our global analysis of changes in chromatin accessibility and nascent transcription nearby annotated human TF motifs, found that changes in response to heat shock were highly correlated between the two cell lines, but that the degree to which some transcription factors were modulated in response to heat shock differed between the two cell lines. These results indicate that the presence of an extra chromosome 21 copy does not disrupt any major signaling events required for the appropriate transcription response immediately following exposure to heat shock stress.

In this study the trisomic cells responded more aggressively to the short heat shock treatment. After heat shocking the lymphoblastoid cell lines for just one hour at 42 °C, we found a more robust activation of HSF1 activity in trisomic cells as inferred from changes in ATAC-seq data, PRO-seq data, and HSF1 regulated steady state transcript levels in scRNA-seq data. The lack of any major heat shock response machinery on chromosome 21 is one reason why heat shock was chosen for this study: we wanted to understand how an extra chromosome might impact global gene regulation as cells mount a transcription program primarily involving genes not present on the aneuploid chromosome. Three genes with links to the heat shock response are present on chromosome 21: HSPA13, DNAJC28 and HSF2BP. It is possible that one of these genes may be impacting the heat shock response directly(Fig ***Figure 1***B). However, the low level expression of these genes in lymphoblastoids and lack of known signaling or transcription factor activities would make a causal role of these genes in increased heat response a surprising mechanism.

HSF1 activation is generally thought of as a response to heat shock but can also occur as a result of heat shock independent stresses such as proteotoxic stress from ribosomal gene imbalances (***Albert et al. (2019***)). Crosstalk from other cell stress pathways like those activated in response to oxidative stress, or ER stress, could also play a part in priming cells to over-respond to heat shock. Chromosome 21 encoded genes such as SOD1, DYRK1A, and four interferon response receptors are directly involved in other cellular stress response pathways. Previous studies have suggested that trisomy caused increased dosage of these genes may lead to the over-activation of stress response pathways in cell and tissue samples from individuals with trisomy 21 (***Sullivan et al. (2016***); ***Araya et al. (2019***); ***Powers et al. (2019***); ***Waugh et al. (2019***); ***Lanzillotta and Di Domenico (2021)***; ***Zhu et al. (2019***); ***Pagano and Castello (2012)***. We do see some evidence, that the interferon response may be overactivated in the trisomic lymphoblastoid cell line in our data Fig ***Figure 3–Figure Supplement 1C***). If any of these other stress response pathways are chronically overactivated in the trisomic cells, crosstalk between these stress responses may cause the overactivated heat shock response observed in this study, and might impact other cellular responses to perturbations.

Despite observing increased heat shock induced HSF1 activity in the trisomic cells, we did not detect a difference between the two cell lines in the transcription levels of the HSF1 transcript itself by PRO-seq or the level of HSF1 transcripts by scRNA-seq ***Figure 5 Figure 5–Figure Supplement 1***. However, HSF1 is a highly regulated TF (***Anckar and Sistonen (2011)***; ***Björk and Sistonen (2010)***; ***Voellmy (2004)***; ***Vihervaara and Sistonen (2014)***), so there are many possible mechanisms that could lead to an elevated HSF1 activation in trisomic cells downstream of HSF1 transcript level changes. A few genes of many with known abilities to regulate HSF1 activity include DAXX, TPR, Mediator, and ribosomal components (***Björk and Sistonen (2010)***; ***Voellmy (2004)***; ***Albert et al. (2019***)). The transcript levels for some of these heat shock regulating factors are elevated in the trisomic cells in our scRNA-seq data ***Figure 5–Figure Supplement 1***. Furthermore, we found that many of these genes were detected in higher quantities in various blood cell types from individuals with trisomy compared to controls, though the results were often inconsistent between blood cell types ***Figure 2***. Note, none of these known HSF1 regulating factors is encoded on chromosome 21, so any trisomy caused misregulation of these genes is likely to be more complex than a trisomy caused gene dosage imbalance. We hypothesize that one of the direct or indirect trisomy caused gene expression changes described above may lead to the observed changes in the regulation of one or more HSF1 regulating genes resulting in an overactivated heat shock response in trisomic blood cells.

Previous studies on the effect of trisomy 21 on stress responses have often focused on stresses known to impact cellular networks regulated by genes on chromosome 21, in which case the increased gene dosage of regulatory genes is expected to lead to a misregulated cellular response. Additionally, measuring the levels of stress response relevant genes in clinical samples can be complicated by the presence of a host of co-morbidity conditions associated with increased occurrence in patients with Down syndrome, but which might be expected to lead to elevated bodily stress. The observation here that a lymphoblastoid cell line with trisomy 21 over-responded to a stress not known to be regulated by a chromosome 21 gene suggests that trisomy 21 may be causing more widespread primary effects on basic gene expression regulation in blood cells than expected.

Because this study began with only a single set of disomic/trisomic cell lines, the next steps for this work need to include more extensive clinical analyses including studying whether this misregulation of HSF1 is a hallmark of Down syndrome and identifying the cell types affected and whether this leads to the depletion or death of any specific immune cell subtype. There are a number of potential consequences that might unfold as a result of misregulating the transcriptional response immediately after heat shock. The irregular blood cell response to heat shock might hamper the ability of the immune system to respond to infection in the presence of a fever in individuals with Down syndrome. Further, if blood cells with Trisomy 21 over-respond to many common cellular stresses, that might mean that a common trisomy associated mechanism may be hampering the ability of patient’s immune systems to respond typically and appropriately to perturbations, and could prove treatable if identified. Future studies would need to investigate whether trisomic blood cells reveal abnormalities in cell survival, or immune cell activation during heat shock stress.

## Acknowledgments

We would like to thank the Translational Nexus Biobank (COMIRB 08-1276), University of Colorado School of Medicine, JFK Partners for access to the cell lines. We acknowledge the BioFrontiers Computing Center at the University of Colorado Boulder for providing High Performance Computing resources supported by BioFrontiers Information Technology. We would also like to thank the BioFrontiers Information Technology for there generous support. We would like to thank the lab of Rui Yi lab, and specifically Dr. Dongmei Wang, for the use of their 10X instrument and their support in producing the scRNA-seq libraries. We would like to acknowledge funding from the Sie Foundation for funding JFC and MAA, and for funding from the 1R01HL156475 MAA and JW, and R01 GM125871 for funding MAA and RDD.

---

**Figure 2–Figure supplement 1.**
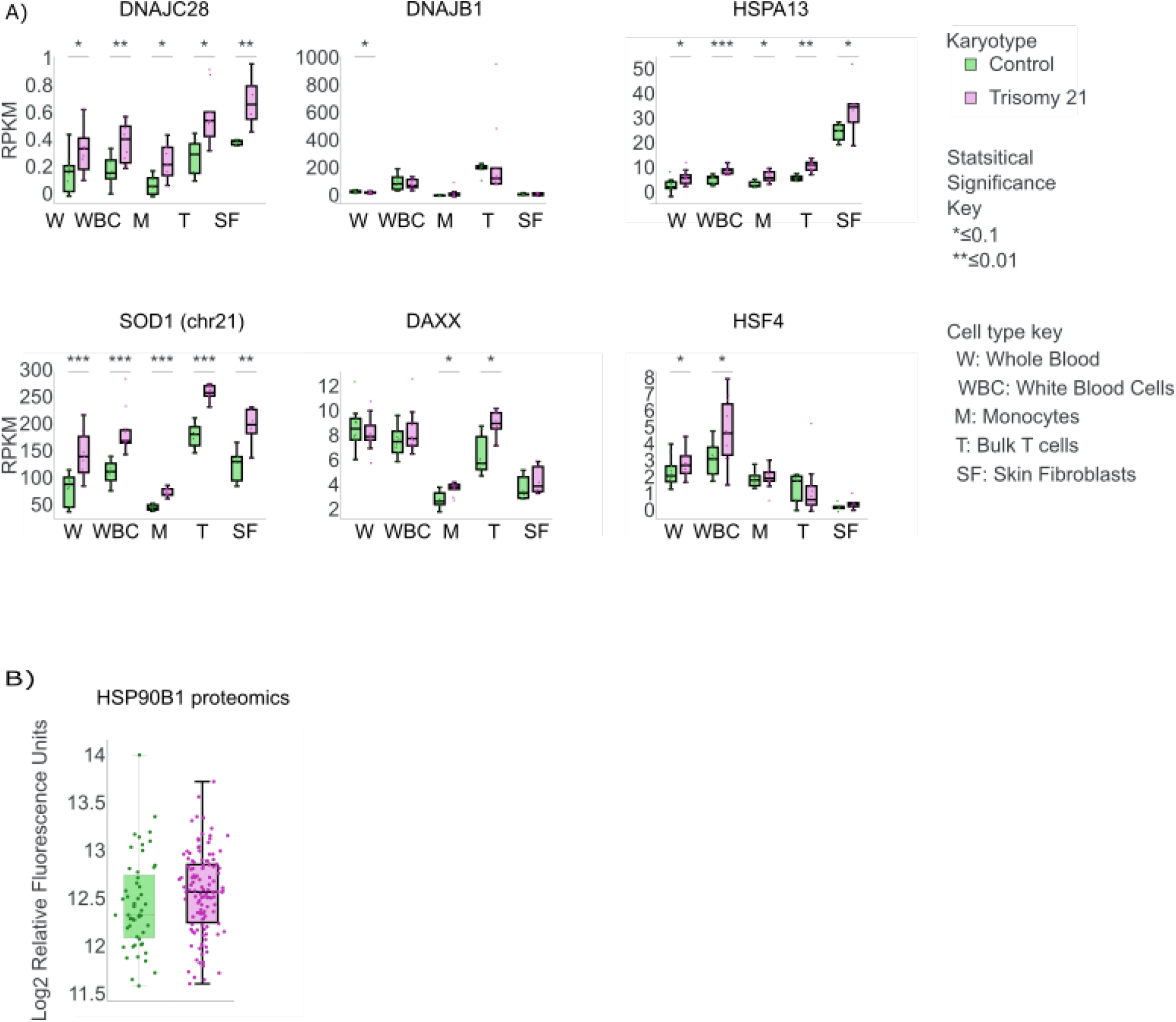
All Data from the Human Trisome Project. A: Heat shock relevant genes (DNAJB1, DNAJC28, HSPA13, SOD1, DAXX, HSF4), are elevated (RNA-seq) in individuals with trisomy 21 (green) relative to disomic controls (green). B: Proteomic data for HSPB1 which shows an increased protein levels in individuals with trisomy 21.

**Figure 3–Figure supplement 1.**
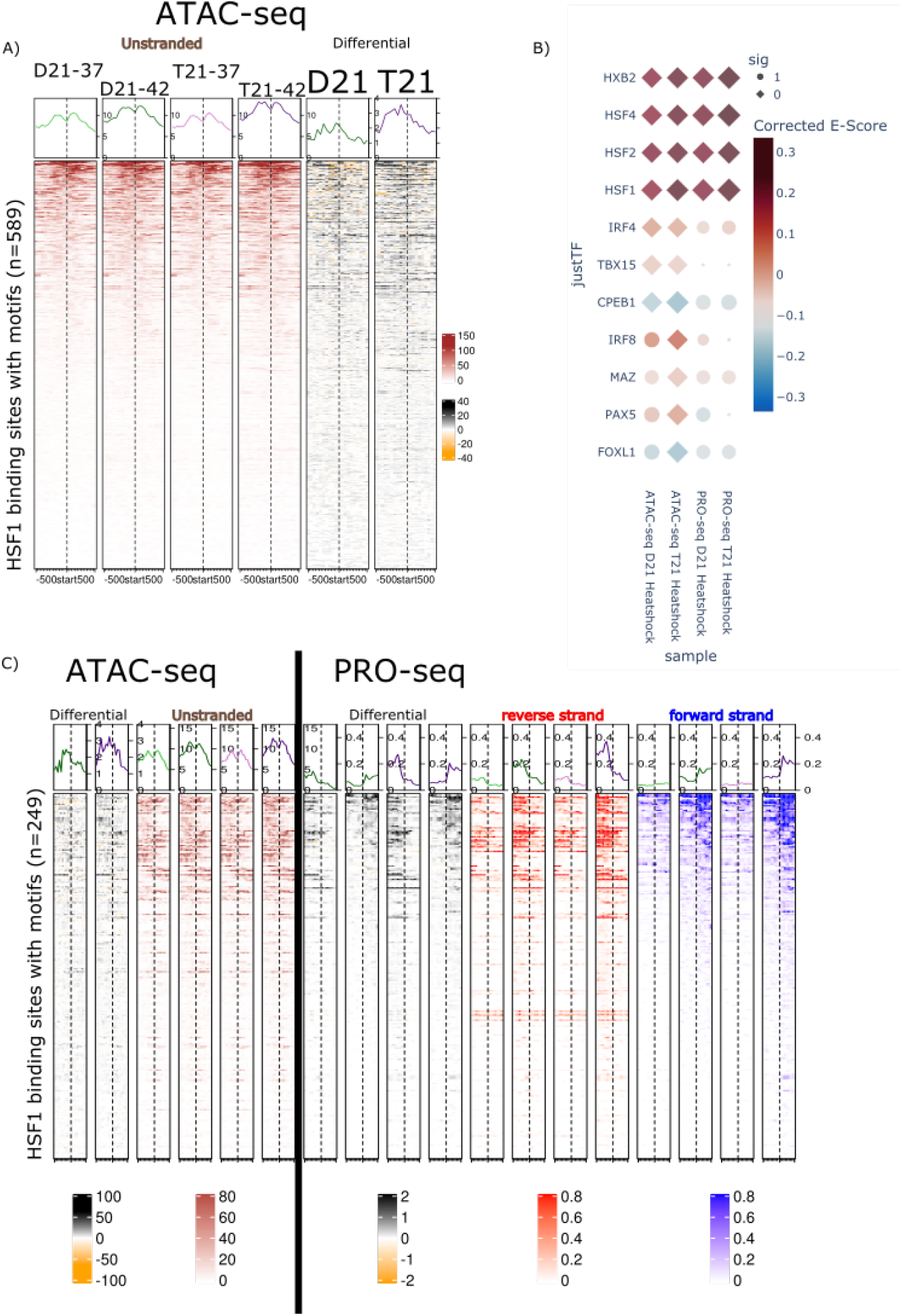
A: Heatmap of ATAC-seq data surrounding HSF1 motifs in HSF1 ChIPseq peaks***Consortium et al. (2012)***. First four columns (in red) show DESeq2 size factor normalized ATAC-seq data surrounding (± 500 nts) a HSF1 motif (dashed center line) at all 589 HSF1 ChIP-seq peaks. Top: line graph of median ATAC-seq depth per position. Fifth and Sixth columns (in black) show the difference (control vs heat shock) in ATAC-seq signal after heat shock. B: A plot showing TFs reported as changed by TFEA in at least one comparison. Diamonds: statically significant (p-value 1×10^−10^), Circles: not significant. Left two columns show the TFEA Escore for ATAC-seq after heat shock in Disomic and Trisomic cell line, whereas the right two columns show TFEA Escore for PRO-seq. Escore, or enrichment score, measures the motif occurrence with open chromatin (ATAC) or transcription (PRO). Hence red Escores indicate enrichment of the motif within regions of increased transcription/accessibility after heatshock whereas blue indicate enrichment of motif within regions of reduced transcription/accessibility. C: Heatmaps of inter-genic HSF1 bound regions for ATAC-seq and PRO-seq data. Columns correspond to: differential ATAC-seq (black, columns 1-2), ATAC-seq signal (red, columns 3-6), differential PRO-seq signal (black, columns 7-10), reverse strand PRO-seq (red, columns 11-14), forward strand PRO-seq (blue, columns 16-18). Top: line graph of median depth per position, dashed line is site of HSF1 motif.

**Figure 4–Figure supplement 1.**
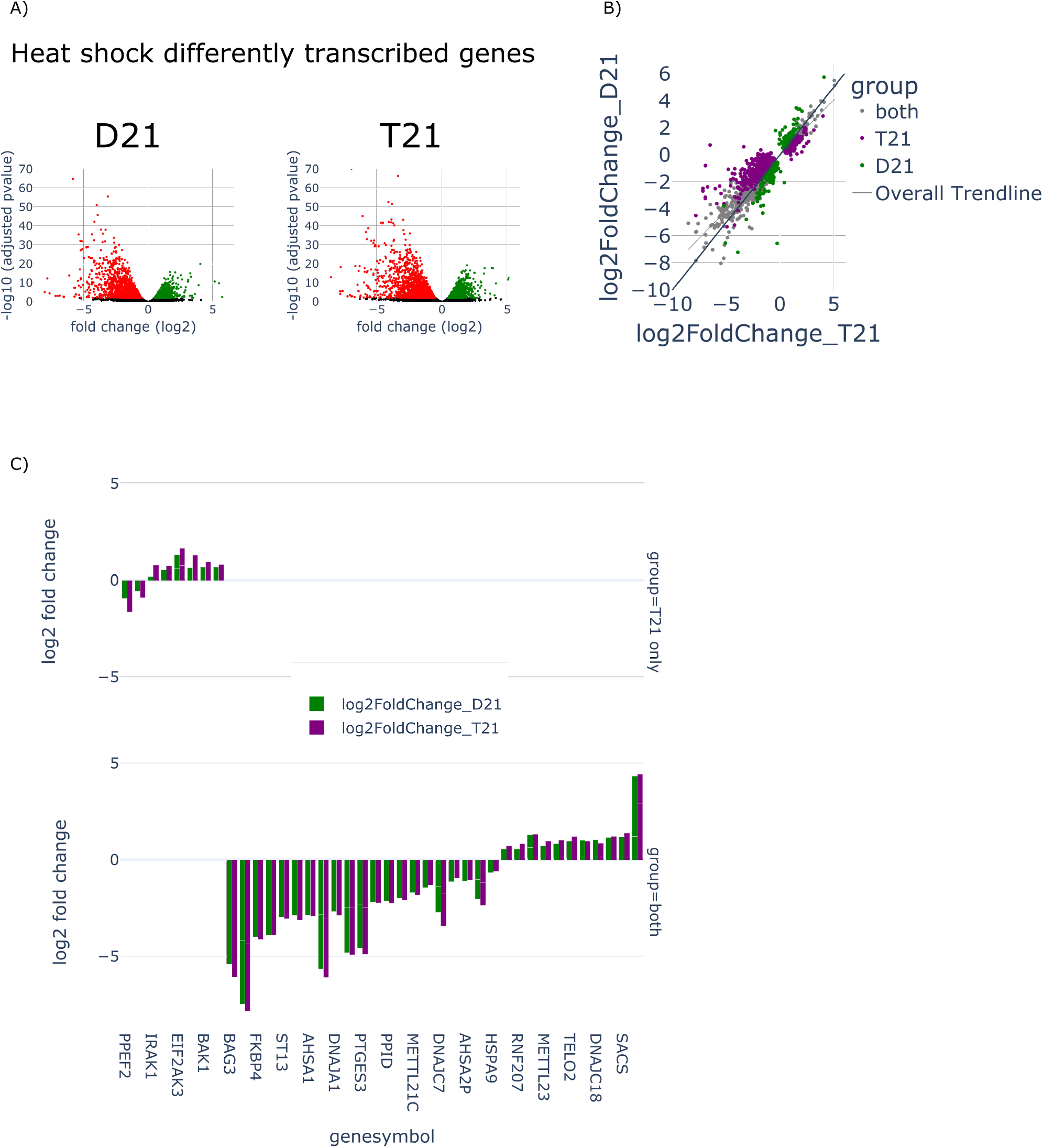
A: Volcano plots of differentially transcribed genes after heat shock. Left: Disomy, Right: Trisomy. Green: up regulated after heat shock, Red: down regulated. B: Scatter plot of the log fold change of the genes changed via heat shock in either trisomy 21 (purple), disomy 21 (green), or both (grey). Best fit line is drawn relative to grey dots only. C: A bar graph of the log fold change of Heat shock genes (as defined by the GO term HEAT_SHOCK_PROTEIN_BINDING). The top plot contains the heat shock genes changed only in the trisomic sample. The bottom plot contains heat shock genes that are differently expressed in both cell types.

**Figure 5–Figure supplement 1.**
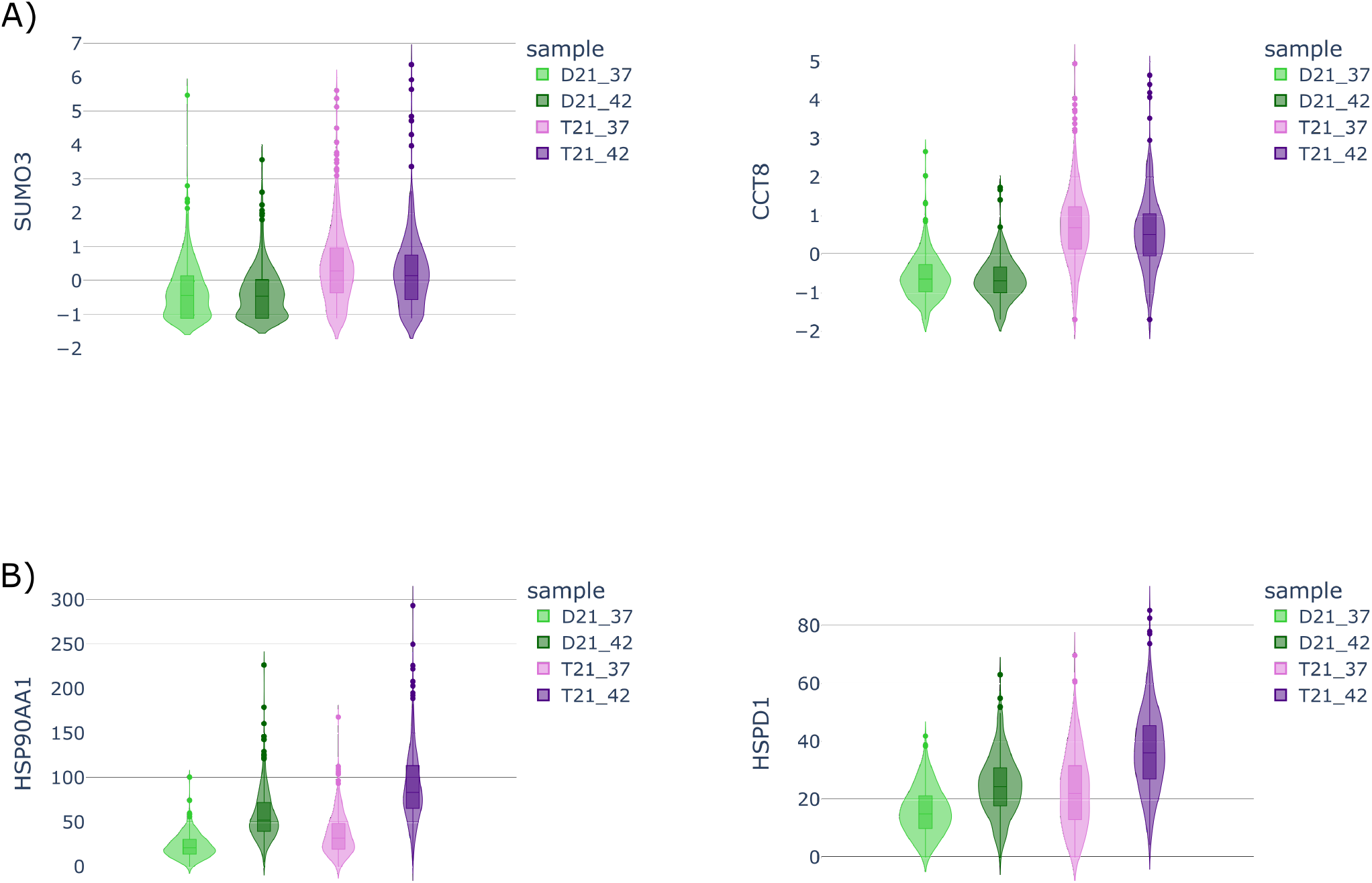
A: The same genes as in **Figure 5**A but with Z scores instead of raw counts. B: The same genes as in **Figure 5**C but with raw counts instead of Z scores.

## References

Aivazidis S, Coughlan CM, Rauniyar AK, Jiang H, Liggett LA, Maclean KN, Roede JR. The burden of trisomy 21 disrupts the proteostasis network in Down syndrome. PloS one. 2017; 12(4):e0176307.

Albert B, Kos-Braun IC, Henras AK, Dez C, Rueda MP, Zhang X, Gadal O, Kos M, Shore D. A ribosome assembly stress response regulates transcription to maintain proteome homeostasis. Elife. 2019; 8:e45002.

Anckar J, Sistonen L. Regulation of HSF1 function in the heat stress response: implications in aging and disease. Annual review of biochemistry. 2011; 80:1089–1115.

Araya P, Waugh KA, Sullivan KD, Núñez NG, Roselli E, Smith KP, Granrath RE, Rachubinski AL, Estrada BE, Butcher ET, et al. Trisomy 21 dysregulates T cell lineages toward an autoimmunity-prone state associated with interferon hyperactivity. Proceedings of the National Academy of Sciences. 2019; 116(48):24231–24241.

Azofeifa JG, Dowell RD. A generative model for the behavior of RNA polymerase. Bioinformatics. 2016 09; 33(2):227–234. doi: 10.1093/bioinformatics/btw599.

Beach RR, Ricci-Tam C, Brennan CM, Moomau CA, Hsu Ph, Hua B, Silberman RE, Springer M, Amon A. Aneuploidy causes non-genetic individuality. Cell. 2017; 169(2):229–242.

Björk JK, Sistonen L. Regulation of the members of the mammalian heat shock factor family. The FEBS journal. 2010; 277(20):4126–4139.

Consortium EP, et al. An integrated encyclopedia of DNA elements in the human genome. Nature. 2012; 489(7414):57.

Huang C, Wu J, Xu L, Wang J, Chen Z, Yang R. Regulation of HSF1 protein stabilization: An updated review. European journal of pharmacology. 2018; 822:69–77.

Lanzillotta C, Di Domenico F. Stress Responses in Down Syndrome Neurodegeneration: State of the Art and Therapeutic Molecules. Biomolecules. 2021; 11(2):266.

Linda Crnic Institute, Human Trisome Project. Global Down Syndrome Foundation; 2021.https://www.trisome.org/.

Mahat DB, Salamanca HH, Duarte FM, Danko CG, Lis JT. Mammalian heat shock response and mechanisms underlying its genome-wide transcriptional regulation. Molecular cell. 2016; 62(1):63–78.

Mei S, Qin Q, Wu Q, Sun H, Zheng R, Zang C, Zhu M, Wu J, Shi X, Taing L, et al. Cistrome Data Browser: a data portal for ChIP-Seq and chromatin accessibility data in human and mouse. Nucleic acids research. 2016; p. gkw983.

Pagano G, Castello G. Oxidative stress and mitochondrial dysfunction in Down syndrome. Neurodegenerative Diseases. 2012; p. 291–299.

Powers RK, Culp-Hill R, Ludwig MP, Smith KP, Waugh KA, Minter R, Tuttle KD, Lewis HC, Rachubinski AL, Granrath RE, et al. Trisomy 21 activates the kynurenine pathway via increased dosage of interferon receptors. Nature communications. 2019; 10(1):1–11.

Ray J, Munn PR, Vihervaara A, Lewis JJ, Ozer A, Danko CG, Lis JT. Chromatin conformation remains stable upon extensive transcriptional changes driven by heat shock. Proceedings of the National Academy of Sciences. 2019; 116(39):19431–19439.

Rubin JD, Stanley JT, Sigauke RF, Levandowski CB, Maas ZL, Westfall J, Taatjes DJ, Dowell RD. Transcription factor enrichment analysis (TFEA): Quantifying the activity of hundreds of transcription factors from a single experiment. Nature Communications Biology. 2021; doi: 10.1038/s42003-021-02153-7.

Sullivan KD, Evans D, Pandey A, Hraha TH, Smith KP, Markham N, Rachubinski AL, Wolter-Warmerdam K, Hickey F, Espinosa JM, Blumenthal T. Trisomy 21 causes changes in the circulating proteome indicative of chronic autoinflammation. Scientific Reports. 2017; 7(1):14818.

Sullivan KD, Lewis HC, Hill AA, Pandey A, Jackson LP, Cabral JM, Smith KP, Liggett LA, Gomez EB, Galbraith MD, DeGregori J, Espinosa JM. Trisomy 21 consistently activates the interferon response. eLife. 2016 jul; 5:e16220.

Tarbell ED, Liu T. HMMRATAC: a Hidden Markov ModeleR for ATAC-seq. Nucleic acids research. 2019; 47(16):e91–e91.

Vihervaara A, Sistonen L. HSF1 at a glance. Journal of cell science. 2014; 127(2):261–266.

Voellmy R. On mechanisms that control heat shock transcription factor activity in metazoan cells. Cell stress & chaperones. 2004; 9(2):122.

Waugh KA, Araya P, Pandey A, Jordan KR, Smith KP, Granrath RE, Khanal S, Butcher ET, Estrada BE, Rachubinski AL, et al. Mass cytometry reveals global immune remodeling with multi-lineage hypersensitivity to type I interferon in Down syndrome. Cell reports. 2019; 29(7):1893–1908.

Zheng R, Wan C, Mei S, Qin Q, Wu Q, Sun H, Chen CH, Brown M, Zhang X, Meyer CA, Liu XS. Cistrome Data Browser: expanded datasets and new tools for gene regulatory analysis. Nucleic Acids Research. 2018 11; 47(D1):D729–D735.

Zhu PJ, Khatiwada S, Cui Y, Reineke LC, Dooling SW, Kim JJ, Li W, Walter P, Costa-Mattioli M. Activation of the ISR mediates the behavioral and neurophysiological abnormalities in Down syndrome. Science. 2019; 366(6467):843–849.

